# Time to first cigarette and its impact on lung tumorigenesis

**DOI:** 10.1101/2022.11.07.515434

**Authors:** Tongwu Zhang, Jian Sang, Neil Caporaso, Fangyi Gu, Amy Hutchinson, Dario Consonni, Angela C. Pesatori, Robert Homer, Stephen Chanock, Maria Teresa Landi

**Author notes:** Corresponding author: Maria Teresa Landi.

## Abstract

Time to first cigarette (TTFC) in the morning has been identified as the best indicator of nicotine dependence and is associated with lung cancer risk beyond other measures of tobacco smoking. Using deep whole genome sequencing of 218 lung cancers from smokers, we show that TTFC is the strongest marker of tobacco smoking mutagenicity, with impact on lung tumor mutational burden, mutational signatures, intratumor heterogeneity, *KRAS* mutations and overall survival. These results pave the way for using TTFC as an easily measurable marker of lung tumorigenesis, with plausible therapeutic and prognostic implications.

## Main

Cigarettes are designed to deliver nicotine to the brain within seconds^1^, making it easier to become dependent on nicotine and more difficult to quit. For this reason, the U.S. Food and Drug Administration (FDA) announced on June 21, 2022, a plan for a proposed rule to establish a maximum level of nicotine in cigarettes. Addition to tobacco smoking has been widely assessed through the Fagerstrom Test for Nicotine Dependence^2^. Among the six items in the test, we^3^ and others^4–6^ have identified the first item, time to first cigarette (TTFC) after waking, as a strong risk factor for lung cancer, independent of other measures of tobacco smoking exposure, i.e., number of cigarettes per day, duration of tobacco smoking in years, age at smoking initiation, and others. TTFC has been considered as a single-item measure of nicotine dependence^7^, biological uptake of nicotine^8^, and smoking cessation success^9^. TTFC has a dose-dependent relationship with the tobacco-specific carcinogen urinary NNAL (4-methylnitrosamino-1-3-pyridyl-1-butanol)^10^. As such, TTFC has been proposed to be a safe, convenient tool for risk stratification to identify smokers who can benefit from lung cancer screening^11^.

We conducted paired whole-genome sequencing analysis (76.4X tumor, 32.6X normal tissue) of 218 lung cancers from smokers (159 adenocarcinomas, 36 squamous cell carcinomas and 23 other histology types) in the Environment and Genetics in Lung cancer Etiology (EAGLE) study^12^ to investigate the impact of different measures of tobacco smoking exposure on lung cancer genomic changes. Multivariate analysis revealed that TTFC was the strongest tobacco smoking variable associated with tumor mutational burden (TMB) and known tobacco smoking mutational signatures (SBS4, DBS2, ID3) (**Fig. 1a**) beyond the number of cigarettes/day, tobacco smoking duration, pack-years of cigarettes, time since last quitting smoking, age at smoking initiation as well as age, sex, histology type, and tumor purity. Shorter TTFC was associated with a higher subclonal mutation ratio indicating higher intratumor heterogeneity (ITH) (P_trend_=0.015). Notably, there is a suggestive association between shorter TTFC and *KRAS* mutations (P_trend_=0.087) that requires follow-up. These findings provide new insights into tobacco-related mutagenesis in lung cancer.

**Fig. 1.**
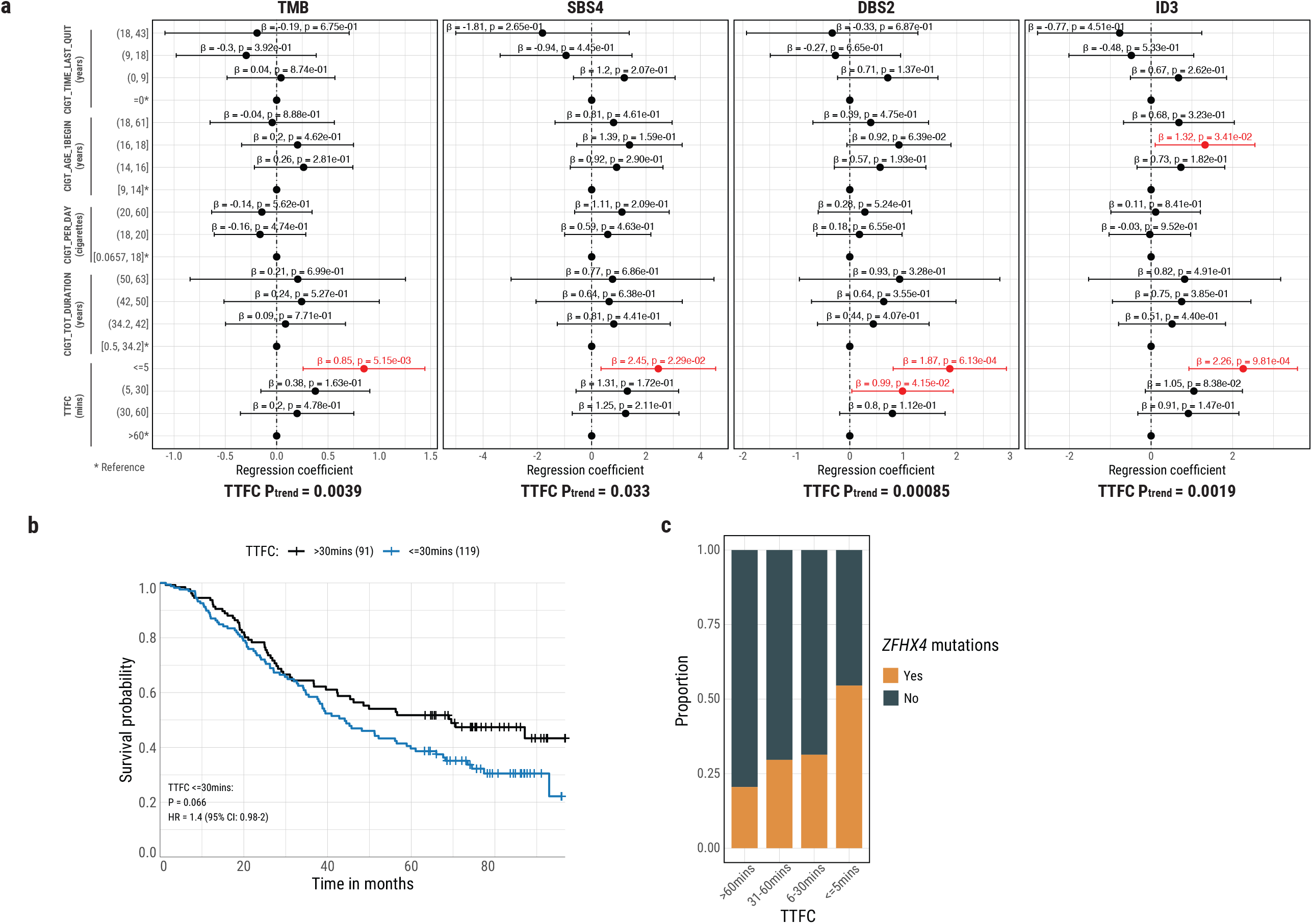
Time to first cigarette (TTFC) and its impact on lung tumorigenesis. **a**) Multivariate regression analysis identified TTFC as the strongest smoking variable associated with TMB, as well as mutations assigned to tobacco smoking-associated signatures (SBS4, DBS2, and ID3). The x-axis shows the regression coefficients, and the y-axis displays the independent smoking variables by category. Significant associations are highlighted in red. The p-values from the trend regression tests (P_trend_) can be found on the bottom of the forest plots. TTFC was the only significant smoking variable with P_trend_ < 0.05. **b**) Kaplan-Meier survival curves for overall survival stratified by TTFC. P-values for significance and hazard ratios (HR) of difference are calculated using the cox proportional hazards model with adjustment for age, sex, stage, histology, and *TP53* mutations. **c)** Proportion plot shows the association between TTFC and nonsynonymous mutation status of *ZFHX4*.

Because of its association with tumor mutational burden^13^, intratumor heterogeneity^14^, and *KRAS* mutation frequency^15^, TTFC could be used to inform lung cancer treatment strategies. We also tested the impact of TTFC on overall survival, after taking into account strong prognostic factors such as age, sex, stage, histology, and *TP53* mutations. Shorter TTFC was associated with poorer survival (HR = 1.4) (**Fig. 1b**).

Mechanistically, shorter TTFC effects could be due to “compensatory smoking”, a tendency to inhale cigarette smoke more deeply in the morning to quickly reach the needed nicotine levels after cessation of smoking during the night^3^. The sudden, intense impact of carcinogens on the lung epithelia can result in high DNA damage that could challenge DNA repair capacity. Further investigations with large-scale sequencing studies and smoking information including TTFC and inhalation habits are required to understand the mutational consequence of TTFC effect through tobacco smoking inhalation.

While we encourage future genomic studies to include data on TTFC, we are aware that most studies do not currently have these data. In a multivariate model, we found that mutations in *ZFHX4* (Zinc finger homeobox protein 4) was a strong marker of shorter TTFC (P_trend_=0.0016; **Fig. 1c**). Mutation frequency of this gene could be used in genomic studies to identify subjects with high nicotine dependence.

In summary, TTFC, an easily measurable tool, can be a surrogate marker of strong mutagenic events in lung tumorigenesis, with plausible therapeutic and prognostic implications. This study emphasizes the importance of the F.D.A. plans to reduce addictiveness of cigarettes and other combusted tobacco products to reduce youth use, addiction, and death.

## Methods

### Assessment of Time to First Cigarette in the morning

Time to first cigarette (TTFC) is the first question of the Fagerstrom Test for Nicotine Dependence. All EAGLE study subjects were asked this question: *“How soon after you wake up do you smoke your first cigarette?”*. The measure was then categorized as <=5 minutes, 5-30 minutes, 30-60 minutes, or >60 minutes.

### Sample collection and whole-genome sequencing (WGS)

We collected lung tumor tissue samples from 218 smokers in The Environment and Genetics in Lung Cancer Etiology (EAGLE) study^12^. Frozen tumor tissues with matched normal control (188 blood, 3 buccal cells, 1 saliva, and 26 normal tissue) samples were used for whole-genome sequencing. DNA extraction and sequencing library construction were performed based on a previously reported protocol^16^. 150bp paired-end whole genome sequencing was performed using the Illumina HiSeq X system by the Broad Institute. The average sequencing depths for tumor and matched control were 76.4x and 32.6x, respectively.

### Bioinformatics analyses of WGS data

The bioinformatics pipelines (https://github.com/xtmgah/Sherlock-Lung) for WGS data processing were similar to those used in our previous publication^16^. Briefly, the raw sequencing data were aligned to the human reference genome GRCh38 following the GATK best practices workflow. Somatic mutations were called using four different algorithms, including MuTect1 (TNsnv in Sentieon), MuTect2 (TNhaplotyper in Sentieon), Strelka, and Sentieon TNscope. Similarly, an ensemble method was applied to merge different callers with additional filtering to reduce false positive calling. Bayesian Dirichlet Process (DPClust)^17–19^ was applied for clustering of subclonal somatic mutations and calculation of subclonal mutation ratio. The Battenberg algorithm^18^ was used for somatic copy number analysis including tumor purity and ploidy estimations. Mutational signatures analyses were performed by the updated computational framework SigProfilerExtractor^20^, based on the SBS288, DBS78, and ID83 profiles. The COSMIC mutation signatures (v3.2) were used as the reference signatures for the known signature deconvolution.

### Statistical and survival analysis

All statistical analyses were performed using the R software v4.1.2. To investigate the association between the mutational landscapes and tobacco smoking variables, we selected 5 independent variables, including TTFC (time to the first cigarette in the morning), CIG_TOT_DURATION (total period (in years) in which the subject smoked cigarettes regularly), CIG_PER_DAY (average intensity of cigarette smoking as number of cigarettes per day), CIGT_AGE_1BEGIN (age when the subject started smoking cigarettes for the first time regularly), and CIGT_TIME_LAST_QUIT (years since quitting smoking cigarettes). Multivariate regression analyses were performed between these smoking variables as categorical values and genomic landscapes including TMB, SBS4, DBS2, ID3, nonsynonymous mutation frequency, and subclonal mutational ratio. The regression model was adjusted for age, sex, histology, and tumor purity. In addition, similar multivariate regression analyses (trend testing) were performed by transforming the smoking variables from categorical variables into continuous variables. For the survival analyses, a proportional-hazards model was used to investigate the associations between the smoking variable TTFC and overall survival, adjusting for age, sex, stage, histology, and *TP53* mutation status.

## Data availability

The 218 normal and tumor paired raw data (BAM files) of the WGS datasets as well as smoking history and other clinical information have been deposited in the dbGaP under accession number phs002992.v1.p1. Researchers will need to obtain authorization from the dbGaP to download these data.

